# The adaptive significance of phasic colony cycles in army ants

**DOI:** 10.1101/091934

**Authors:** Simon Garnier, Daniel J. C. Kronauer

**Affiliations:** Department of Biological Sciences, New Jersey Institute of Technology, Newark, NJ 07102, USA; Laboratory of Social Evolution and Behavior, The Rockefeller University, New York, NY 10065, USA

**Keywords:** Dorylinae, Evolution, Foraging, Formicidae, Group predation, Modelling, Nomadism, Reproductive cycle

## Abstract

Army ants are top arthropod predators in tropical forests around the world. The colonies of many army ant species undergo stereotypical behavioral and reproductive cycles, alternating between brood care and reproductive phases. In the brood care phase, colonies contain a cohort of larvae that are synchronized in their development and have to be fed. In the reproductive phase larvae are absent and oviposition takes place. Despite these colony cycles being a striking feature of army ant biology, their adaptive significance is unclear. Here we use a modelling approach to show that cyclic reproduction is favored under conditions where per capita foraging costs decrease with the number of larvae in a colony ("High Cost of Entry" scenario), while continuous reproduction is favored under conditions where per capita foraging costs increase with the number of larvae ("Resource Exhaustion" scenario). We argue that the former scenario specifically applies to army ants, because large raiding parties are required to overpower prey colonies. However, once raiding is successful it provides abundant food for a large cohort of larvae. The latter scenario, on the other hand, will apply to non-army ants, because in those species local resource depletion will force workers to forage over larger distances to feed large larval cohorts. Our model provides the first quantitative framework for understanding the adaptive value of phasic colony cycles in ants.

## Introduction

Army ants are top arthropod predators in tropical rain forests around the world (Schneirla 1971; Gotwald 1995; Kronauer 2009). Their colonies can measure hundreds of thousands or even millions of individuals in size, and their live prey is overwhelmed by the mass onslaught of large raiding parties, making army ants the ultimate group hunters. Army ant colonies emigrate frequently, a behavior that is thought to be related to the necessity of regularly exploring new hunting grounds after depleting local prey patches (Gotwald 1995; Kronauer 2009; Schöning 2005). While most social insect colonies are founded by a single female (the queen) and then slowly grow to mature size, army ant colonies multiply by a process called colony fission, during which a large colony splits into two roughly equally sized daughter colonies (Schneirla 1971; Gotwald 1995; Kronauer 2009). This unusual mode of reproduction is arguably related to the fact that only large army ant colonies can mount successful raids, while small incipient colonies would be unviable.

The colonies of many army ant species undergo stereotypical behavioral and reproductive cycles, during which colonies alternate between brood care and reproductive phases (Figure 1; Schneirla 1971; Gotwald 1995; Kronauer 2009; Oxley et al. 2014). Colony emigrations are usually restricted to the brood care phase, and foraging activity is highly intensified during that phase. The brood care phase has therefore also been referred to as the “nomadic” or “foraging” phase, and the reproductive phase has been referred to as the “statary” phase (“statary” meaning “settled”) by previous authors (Schneirla 1971; Ravary & Jaisson 2002). This phasic lifestyle has evolved repeatedly across the ant phylogeny, and it can therefore be found in distantly related species. While not all army ants are phasic, all known phasic species have army ant-like biology. Examples of phasic species in the subfamily Dorylinae, which encompasses the vast majority of army ants, include the well-studied Neotropical army ants *Eciton burchellii* and *E. hamatum*, the North American *Neivamyrmex nigrescens*, as well as the Asian *Aenictus laeviceps* (reviewed in Schneirla 1971; Gotwald 1995). Several additional species in the doryline genera *Aenictus, Cerapachys, Cheliomyrmex, Eciton, Neivamyrmex, Nomamyrmex*, and *Sphinctomyrmex* appear to be phasic (Rettenmeyer 1963; Schneirla 1971; Hölldobler 1982; Buschinger et al. 1989; Gotwald 1995). The only ant species in which phasic colony cycles can readily be manipulated experimentally, the clonal raider ant *Ooceraea biroi* (formerly *Cerapachys biroi)*, also belongs to this subfamily (Ravary & Jaisson 2002; Ravary et al. 2006; Teseo et al. 2013; Oxley et al. 2014; Libbrecht et al. 2016; Ulrich et al. 2016). Outside of the Dorylinae, phasic species with army ant-like biology can be found in the subfamilies Ponerinae (genus *Simopelta;* Gotwald & Brown 1966; Kronauer et al. 2011), Leptanillinae (genus *Leptanilla;* Masuko 1990), and Amblyoponinae (genus *Onychomyrmex;* Miyata et al. 2003).

**Figure 1.**
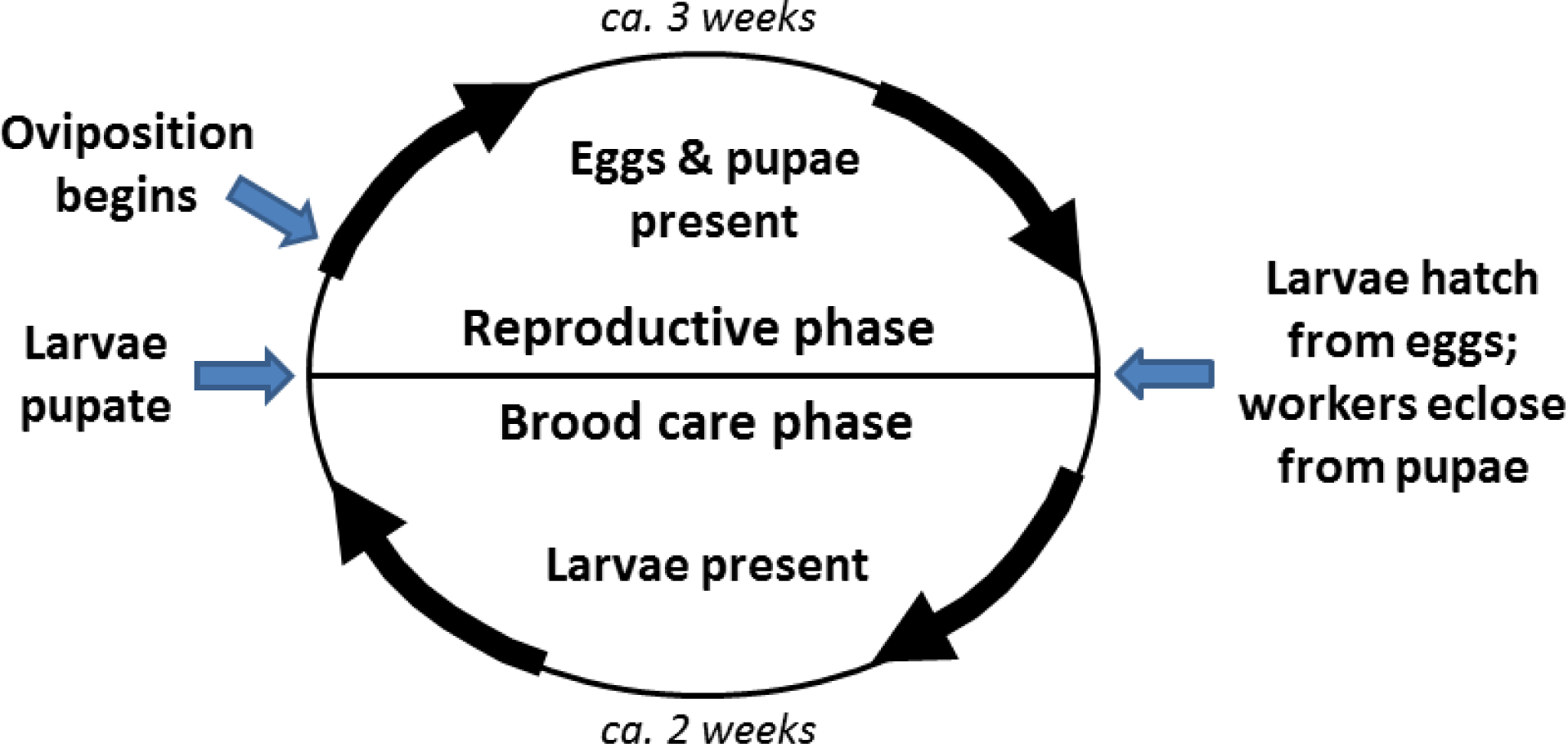
Schematic of a generalized phasic army ant colony cycle. Larvae are present during the brood care phase and develop in discrete cohorts. Larvae have two main effects on adults: they suppress ovarian activity and oviposition, and induce brood care behavior, including nursing and foraging (Ravary et al. 2006; Teseo et al. 2013; Ulrich et al. 2016). As larvae pupate toward the end of the brood care phase the colony transitions to the reproductive phase. Oviposition commences a few days into the reproductive phase. As larvae hatch and young workers emerge from pupae toward the end of the reproductive phase the colony transitions to the next brood care phase. In *Eciton burchellii, E. hamatum*, and *Ooceraea biroi*, the reproductive and brood care phases last for ca. three and two weeks, respectively (Schneirla 1971; Ravary & Jaisson 2002).

The fact that phasic colony cycles have evolved repeatedly in species with army ant-like biology suggests that they represent specific adaptations to the army ant lifestyle. While we are beginning to understand the proximate mechanisms underlying phasic colony cycles in great detail using the clonal raider ant as a laboratory model system (Ravary & Jaisson 2002; Ravary et al. 2006; Teseo et al. 2013; Oxley et al. 2014; Libbrecht et al. 2016; Ulrich et al. 2016), the ultimate adaptive significance of phasic cycles has remained elusive. Arguably the most general hypothesis that has been put forward is that in at least some species with an army ant-like biology phasic colony cycles minimize the overall cost of foraging by temporally restricting the presence of food-demanding larvae (Kronauer 2009). Here we assess the plausibility of this hypothesis by developing an explicit model that integrates alternative reproductive strategies (phasic vs. non-phasic) with the costs associated with different foraging scenarios. In particular, we investigate three possible foraging scenarios: (1) the cost of foraging scales proportionally with the number of larvae to be fed; (2) the cost of foraging increases proportionally faster for smaller numbers of larvae than for larger ones (army ant-like foraging scenario); (3) the cost of foraging increases proportionally slower for smaller numbers of larvae than for larger ones (non-army ant-like foraging scenario). We find that a phasic lifestyle indeed minimizes the likely costs associated with group predation (scenario 2), while a non-phasic lifestyle minimizes the costs associated with other forms of foraging (scenario 3), thereby providing a convincing adaptive scenario for the evolution of army ant colony cycles.

## 2. Methods

### 2.1 Modelling colony reproductive strategy

The relative number of larvae *l* in the colony at time *t* is modeled as a function of the form:

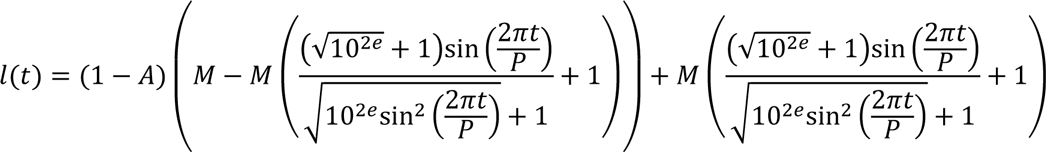

where *M* represents the average of the periodical wave, *P* its period, and *A* its amplitude. The amplitude *A* is relative to the average *M* of the periodical wave. When *A = 1* the minimum value of the wave is 0 and the maximum is 2*M.* When *A =* 0 the wave is flat (i.e. its minimum and maximum values are both equal to the average of the wave). The exponent *e* controls the degree of “squarity” of the wave. Positive values of *e* return a more square-like wave while negative values return a more sine-like wave. This allows us to control how smooth the reproductive cycle is, in other words, how gradual or abrupt the transitions between brood care and reproductive phases are.

For the remainder of this study, we will arbitrarily set the value of *P*, i.e. the length of the reproductive cycle, to 1. We will also set the value of *M*, i.e. the average relative number of larvae in the colony, to 0.5. As a consequence, both the absolute length of the reproductive cycle and the absolute number of larvae a colony raises per reproductive cycle are constant across all comparisons. With *P = 1* and *M =* 0.5 we can then simplify the previous equation as follows:

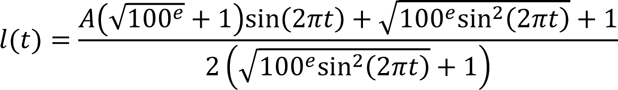

Figure 2 shows the effect of varying either the amplitude *A*, or the “squarity exponent” *e* on the temporal dynamics of the relative number of larvae present in the colony across the reproductive cycle.

**Figure 2.**
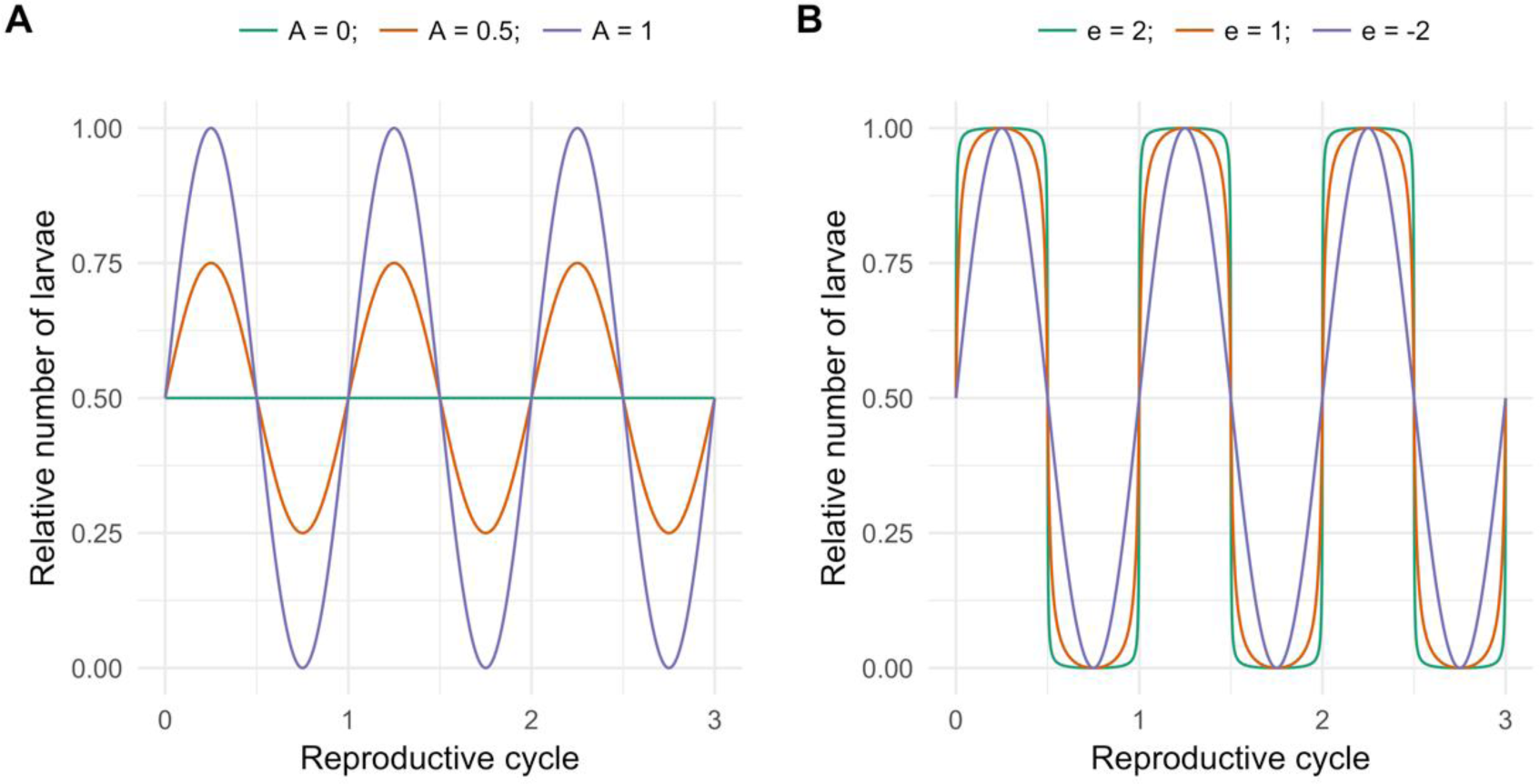
Effect of varying **(A)** the amplitude *A* and **(B)** the “squarity exponent” *e* on the temporal dynamics of the relative number of larvae present in the colony across the reproductive cycle.

### 2.2. Modelling colony foraging cost

We consider three possible scenarios for the distribution of foraging costs as a function of the number of larvae that have to be fed:

1. “Proportional”: In this scenario, the cost of foraging grows linearly with the number of larvae. This scenario is biologically unlikely but will serve as a baseline comparison for the performance of the other two scenarios.
2. “High Cost of Entry”: In this scenario, the cost of foraging increases proportionally faster for smaller numbers of larvae than for larger ones. This corresponds to cases where a minimum number of workers are required before foraging yields significant benefits (for instance where ants have to overpower large prey items or other social insect colonies). This is the scenario likely faced by many ant species with army ant-like biology.
3. “Resource Exhaustion”: In this scenario, the cost of foraging increases proportionally slower for smaller numbers of larvae than for larger ones. This corresponds to cases where local resources are exploited faster than they are replenished, which forces workers to cover increasingly larger foraging distances as the number of larvae increases. This is the scenario that is likely faced by ant species that mainly forage as scavengers, herbivores, or individual predators, i.e. all ant species except those with army ant-like biology.

For all three scenarios, we can model the change in foraging cost *c* as a function of the relative number of larvae *l* with a function of the form:

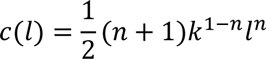

where *k* is the maximum number of larvae that a colony can have at any given time, and *n* is a parameter that determines how the cost of foraging scales with the number of larvae to be fed. When *n = 1*, the cost of foraging scales linearly with the number of larvae ("Proportional" scenario). When *n > 1*, the cost of foraging grows slower for smaller than for larger numbers of larvae ("Resource Exhaustion" scenario). When *0* ≤ *n < 1*, the cost of foraging grows faster for smaller than for larger numbers of larvae ("High Cost of Entry" scenario).

Note that this function is designed to ensure that its integral between 0 and *k* is the same regardless of the value of *n*, hence normalizing the foraging cost between all possible values of *n.*

For the remainder of this study, we will set *k = 1*, which allows us to simplify the previous equation as follows:

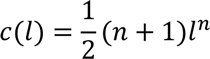

Figure 3 shows the effect of varying *n* on the shape of the foraging cost function.

**Figure 3.**
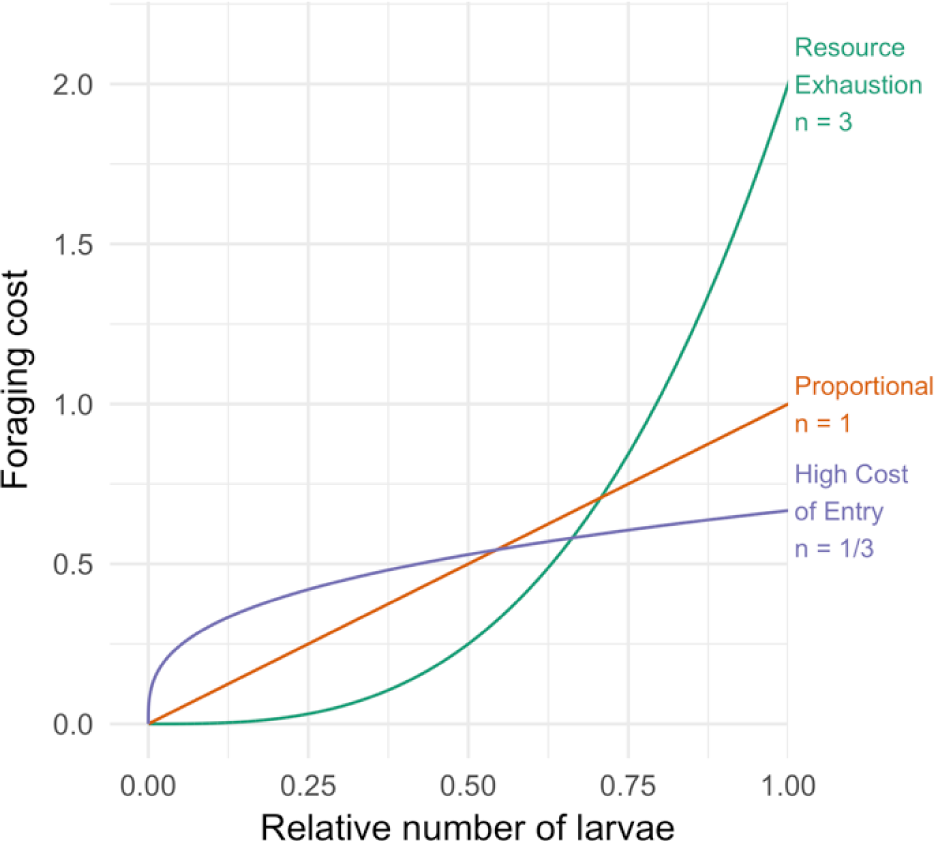
Effect of varying the scaling parameter *n* on the shape of the foraging cost function. The “Proportional” scenario is obtained with *n = 1.* The “High Cost of Entry” scenario is obtained with *0* ≤ *n* < 1 (*n* = 1/3 is shown here as an example). Finally, the “Resource Exhaustion” scenario is obtained with *n* > 1 (*n* = 3 is shown here as an example).

### 2.3. Integrating reproductive strategy and foraging cost

To evaluate the performance of a given reproductive strategy under different foraging cost distributions, we calculate the total foraging cost (i.e. we integrate the composite function *c*(*l*(*t*))) across one entire colony cycle for different values of the relative amplitude *A* of the reproductive cycle, the "squarity exponent" *e* of the reproductive cycle, and the foraging cost scaling parameter *n.* The general shape of the integral function is as follows:

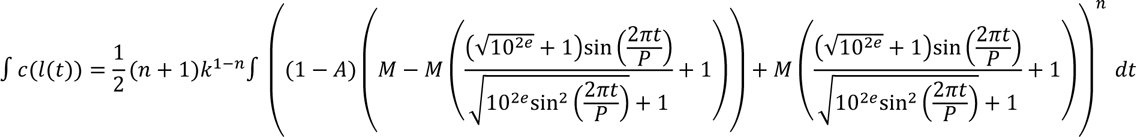

With *P*=1, *M*=0.5 and *k*=1, we can simplify this equation as follows:

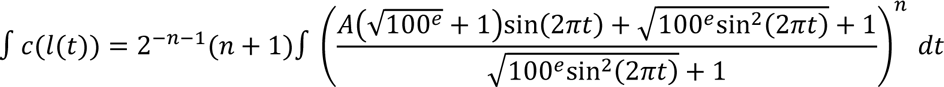

### 2.4. Software

We used Mathematica 11.0.1.0 to simplify the equations and generate the integral of the function combining the reproductive strategy with the foraging cost: 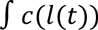.

We used the “integrate” function in the “stats” package (version 3.3.2) in R (version 3.3.2) to calculate the value of the integral 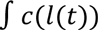 over various values of the parameters *A, e*, and *n.*

All figures were generated in R using the ggplot2 (version 2.2.0) and cowplot (version 0.7.0) packages.

All code used in this manuscript can be found at https://goo.gl/ClHwDz.

## Results

### 3.1. Cost of foraging under different foraging scenarios and reproductive strategies

Using the integral function described above, we compute the total cost of foraging over a colony cycle for various combinations of the relative amplitude *A* of the reproductive cycle and the shape parameter *n* of the foraging cost function. Since we are not interested here in the effect of the shape of the reproductive cycle, which will be treated in Section 3.2, we set its shape to a near-square wave (*e* = 10). Note, however, that the results are qualitatively equivalent with a sine wave. Figure 4 summarizes the results.

**Figure 4.**
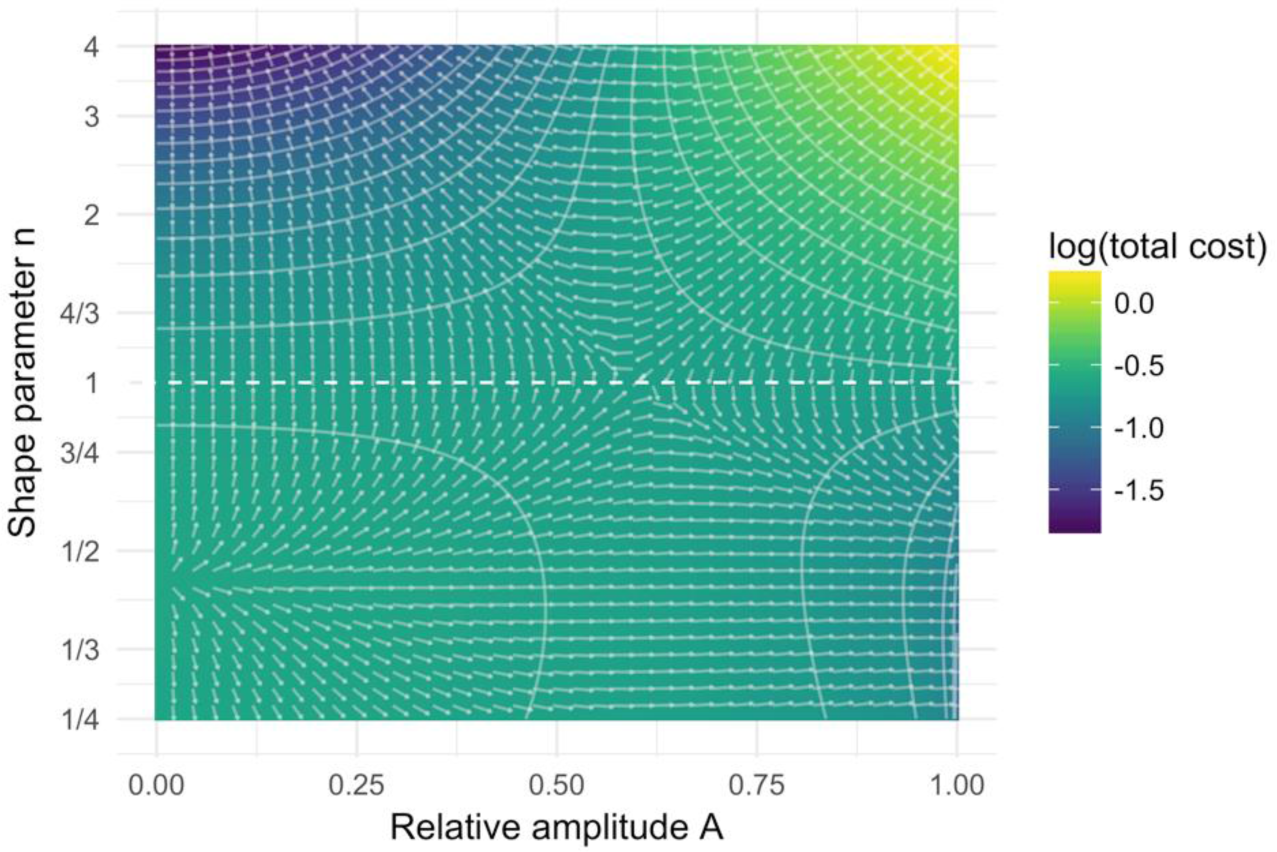
Total foraging cost over a reproductive cycle for various combinations of the relative amplitude *A* of the reproductive cycle and the shape parameter *n* of the foraging cost function. Values of *n* above the dashed line (which indicates the “Proportional” scenario) correspond to “Resource Exhaustion” scenarios, while values below the dashed line correspond to “High Cost of Entry” scenarios. Isolines (white contour lines) represent points in the parameter space with constant value, and gradient vectors (white arrows) represent the direction, but not the intensity of the local gradient.

When the foraging cost function corresponds to a “Resource Exhaustion” scenario (*n >* 1), the lowest total foraging cost is obtained by a non-phasic reproductive strategy that distributes the number of larvae produced by the colony uniformly in time (i.e. *A =* 0; see top left part of Figure 4).

On the contrary, when the foraging cost function corresponds to a “High Cost of Entry” scenario (0 ≤ *n <* 1), the total foraging cost decreases with periodical variation in the number of larvae produced by the colony. Lower costs are achieved for larger oscillation amplitudes, and colonies perform best under the most extreme phasic reproductive strategy in terms of oscillation amplitude (i.e. *A =* 1; see bottom right part of Figure 4).

### 3.2. Effect of smoothness of phase transitions

Here we test the effect of abrupt phase transitions in which all larvae hatch and pupate at the exact same time (i.e. a square wave cycle) versus smooth phase transitions in which larvae hatch and pupate around an average time (i.e. a sine wave cycle) in a “High Cost of Entry” scenario. We do not test this effect in a “Resource Exhaustion” scenario because the results in Section 3.1 show that a perfectly non-phasic reproductive strategy is favored in this case.

Using the integral function described above, we compute the total cost of foraging over a reproductive cycle for different values of the cycle’s “squarity exponent” *e*. We set *A* to 1, i.e. a cycle with maximum oscillation intensity, and *n* to 1/4, i.e. a “High Cost of Entry” scenario. Note that results are qualitatively similar for any combination of *A >* 0 and 0 < *n < 1*. Figure 5 summarizes the results.

**Figure 5.**
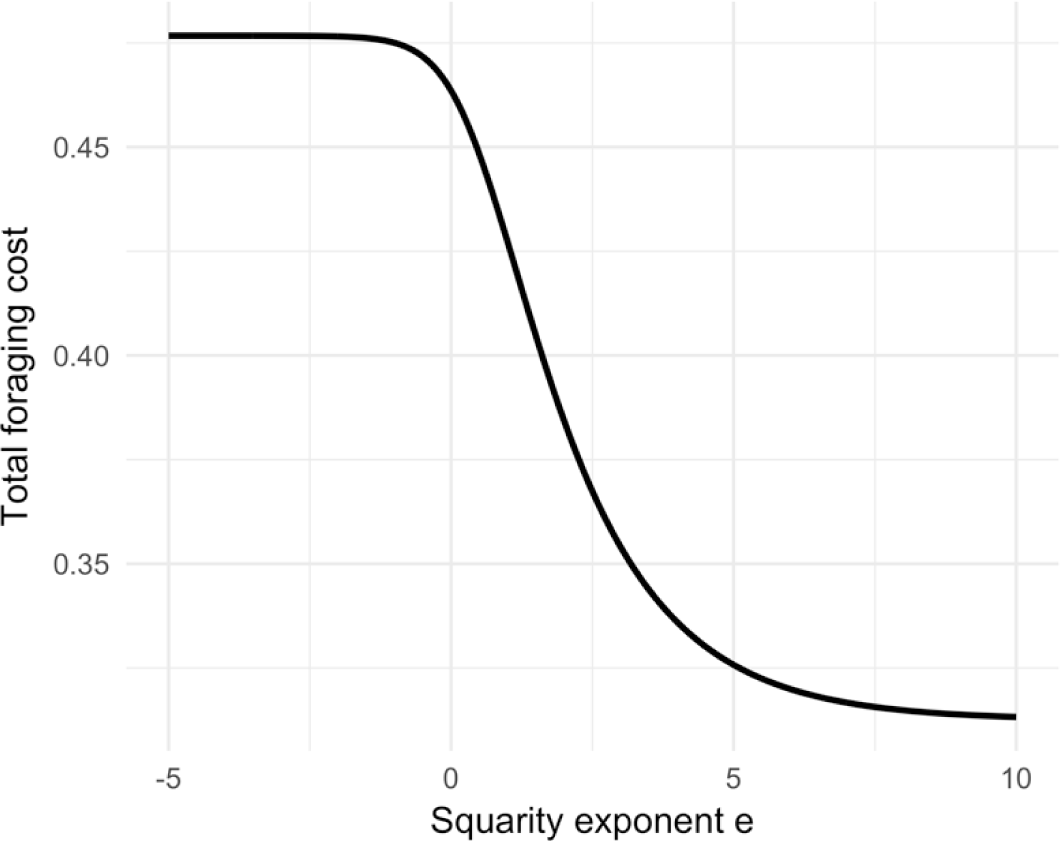
Effect of the smoothness of phase transitions (determined by the “squarity exponent” *e)* on the total foraging cost over a reproductive cycle for colonies in a “High Cost of Entry” foraging scenario.

The total cost of foraging decreases with the value of the “squarity exponent” *e*, indicating that abrupt phase transitions are more beneficial than smooth phase transitions for colonies experiencing a “High Cost of Entry” foraging scenario.

## 4 Discussion

Our model shows that phasic colony cycles are adaptive in species where the relative cost of foraging, that is the investment required per larva, is high when few larvae have to be fed, but decreases as the number of larvae increases (Figure 4). Such a “High Cost of Entry” scenario applies, for example, when a substantial investment into foraging is required before foraging yields any benefits at all. This seems to be the case in many army ants, because large prey items or other social insect colonies can only be overpowered by large raiding parties, while individuals or small groups of foragers will not be successful. In other words, for much of the parameter space an army ant colony should either mount a full-blown attack or not forage at all. A phasic cycle allows army ant colonies to do exactly that: during the brood care phase costly raids bring in lots of food to feed large cohorts of larvae, while little or no foraging is required during the reproductive phase when no larvae have to be fed.

Our model further shows that, under a “High Cost of Entry” scenario, phase transitions should be as abrupt as possible (Figure 5). In other words, the larvae of a given cohort should be perfectly synchronized in their development. While larval development in phasic army ants is indeed tightly synchronized, this synchronization is not perfect, and different larval instars overlap to some extent during the brood care phase (Schneirla 1971; Ravary & Jaisson 2002). However, this overlap could simply reflect a tradeoff between minimizing the period of reproductive activity to maximize developmental synchronization and the incentive to produce large brood cohorts.

Outside of army ants, foraging strategies are extremely diverse across different ant species (Lanan 2014). However, in the vast majority of cases increased investment in foraging is unlikely to yield the same disproportionate returns via synergistic effects of foraging in large groups. In particular, as foraging intensity increases, these species should suffer from “Resource Exhaustion” as the resources in the immediate vicinity of the nest become depleted and foragers have to cover larger distances to encounter food. In other words, the per capita cost of feeding larvae increases with the number of larvae that have to be provided for. Our model shows that these species should therefore be non-phasic, and indeed no phasic species without an army ant-like biology are known.

Interestingly, not all army ants are phasic. Prominent examples of non-phasic army ants include the Old World genus *Dorylus* (Schneirla 1971; Schöning 2005), as well as the New World genus *Labidus* (Rettenmeyer 1963). Even though colonies of non-phasic army ants still emigrate frequently, possibly in response to local food depletion or predator attack, the emigrations do not follow a regular temporal pattern and do not coincide with the presence of particular developmental stages (Rettenmeyer 1963; Schneirla 1971; Schöning 2005). According to our model, we would predict that in these species foraging is more efficient with a relatively small investment. In other words, the relationship between foraging cost and the number of larvae should constitute a “Resource Exhaustion” scenario rather than a “High Cost of Entry” scenario as in phasic army ants (Figure 4). In this context it is interesting to note that both *Dorylus* and *Labidus* colonies are unusually large, containing from over a million to several million workers, while the colonies of other army ants are at least one or two orders of magnitude smaller (Rettenmeyer 1963; Schneirla 1971; Gotwald 1995). It is therefore possible that even raiding parties that are large in absolute size and therefore can forage efficiently come at a small cost at the colony level in these species, because they still represent only a small fraction of the total worker force. Furthermore, while most army ants exclusively or predominantly prey on other social insects, social insects constitute only a small proportion of the prey in *Dorylus* and *Labidus*, whose prey spectra are generally much broader than those of other army ants (Rettenmeyer 1963; Gotwald 1995; Schöning et al. 2008). In fact, *Dorylus* and *Labidus* are the only army ants that also forage on things other than live prey, including animal carcasses, nuts, fruits, grains, and vegetable oil (Gotwald 1995). This implies that food might be more readily available for *Dorylus* and *Labidus* army ants. Furthermore, less relative investment into foraging could still have positive returns because it can be directed toward plant material, animal carcasses, or prey that can be more easily overwhelmed than the well-fortified colonies of social insects. These and possibly other factors might well place non-phasic army ants in a “Resource Exhaustion” scenario. However, given the difficulty of working with army ants in the field, it will be challenging to experimentally quantify the cost of foraging in relation to the number of larvae to be fed for any given species.

## Authors’ contributions

SG and DJCK conceived the study and developed the model. SG implemented and analyzed the model. SG and DJCK wrote the paper.

## Acknowledgements

We thank Yuko Ulrich and Jonathan Saragosti for discussing the ideas underlying this study with us.

## References

Buschinger, A., Peeters, C., Crozier, R.H. (1989). Life-pattern studies on an Australian *Sphinctomyrmex* (Formicidae: Ponerinae: Cerapachyini): functional polygyny, brood periodicity and raiding behaviour. Psyche 96: 287–300.

Gotwald, W.H. Jr, Brown, W.L. Jr (1966). The ant genus *Simopelta* (Hymenoptera: Formicidae). Psyche 73: 261–277.

Gotwald, W.H. Jr. (1995). Army ants: the biology of social predation. Cornell University Press.

Hölldobler, B. (1982). Communication, raiding behaviour and prey storage in *Cerapachys* (Hymenoptera: Formicidae). Psyche 89: 3–23.

Kronauer, D.J.C. (2009). Recent advances in army ant biology (Hymenoptera: Formicidae). Myrmecological News 12: 51–65.

Kronauer, D.J.C., O’Donnell, S., Boomsma, J.J., Pierce, N.E. (2011). Strict monandry in the ponerine army ant genus *Simopelta* suggests that colony size and complexity drive mating system evolution in social insects. Molecular Ecology 20: 420–428.

Lanan, M. (2014). Spatiotemporal resource distribution and foraging strategies of ants (Hymenoptera: Formicidae). Myrmecological News 20: 53–70.

Libbrecht, R., Oxley, P.R., Keller, L., Kronauer, D.J.C. (2016). Robust DNA methylation in the clonal raider ant brain. Current Biology 26: 391–395.

Masuko, K. (1990). Behavior and ecology of the enigmatic ant *Leptanilla japonica* Baroni Urbani (Hymenoptera: Formicidae: Leptanillinae). Insectes Sociaux 37: 31–57.

Miyata, H., Shimamura, T., Hirosawa, S., Higashi, S. (2003). Morphology and phenology of the primitive ponerine army ant *Onychomyrmex hedleyi* (Hymenoptera: Formicidae: Ponerinae) in a highland rainforest of Australia. Journal of Natural History 37: 115–125.

Oxley, P.R., Ji, L., Fetter-Pruneda, I., McKenzie, S.K., Li, C., Hu, H., Zhang, G., Kronauer, D.J.C. (2014). The genome of the clonal raider ant *Cerapachys biroi*. Current Biology 24: 451–458.

Ravary, F., Jaisson, P. (2002). The reproductive cycle of thelytokous colonies of *Cerapchys biroi* Forel (Formicidae, Cerapachyinae). Insectes Sociaux 49: 114–119.

Ravary, F., Jahyny, B., Jaisson, P. (2006). Brood stimulation controls the phasic reproductive cycle of the parthenogenetic ant *Cerapachys biroi*. Insectes Sociaux 53: 20–26.

Rettenmeyer, C.W. (1963). Behavioral studies of army ants. University of Kansas Science Bulletin 44: 281–465.

Schneirla, T.C. (1971). Army ants: a study in social organization. W.H. Freeman & Co.

Schöning, C., Njagi, W.M., Franks, N.R. (2005). Temporal and spatial patterns in the emigrations of the army ant *Dorylus (Anomma) molestus* in the montane forest of Mt Kenya. Ecological Entomology 30: 532–540.

Schöning, C., Njagi, W., Kinuthia, W. (2008). Prey spectra of two swarm-raiding army ant species in East Africa. Journal of Zoology 274: 85–93.

Teseo, S., Kronauer, D.J.C., Jaisson, P., Châline, N. (2013). Enforcement of reproductive synchrony via policing in a clonal ant. Current Biology 23: 328–332.

Ulrich, Y., Burns, D., Libbrecht, R., Kronauer, D.J.C. (2016). Ant larvae regulate worker foraging behavior and ovarian activity in a dose-dependent manner. Behavioral Ecology and Sociobiology 70: 1011–1018.

